# SPP1+ tumor-associated macrophages induce radioresistance of nasopharyngeal carcinoma through crosstalk with the tumor microenvironment

**DOI:** 10.1101/2024.10.23.619837

**Authors:** Wenda Zhang, Lili Lin, Ying Haoxuan, Nan Li, Xinyi Zhou, Qi Yang, Shulu Hu, Jiayi Cai, Anqi Lin, Cheng Quan, Zaoqu Liu, Jian Zhang, Peng Luo, Ting Wei

**Author notes:** Corresponding Authors: mail addresses (Jian Zhang); (Peng Luo); (Ting Wei). Joint Authors: These authors have contributed equally to this work and share first authorship (Wenda Zhang, Lili Lin, Ying Haoxuan, Nan Li).

## Abstract

The function and phenotype of tumor-associated macrophages (TAMs) influence the efficacy of radiotherapy. However, the effect of different subsets of TAMs on the radioresistance in nasopharyngeal carcinoma (NPC) still needs to be further explored. This study utilized single-cell RNA sequencing (scRNA-seq) to analyze 120,579 cells obtained from 15 NPC samples. Cell annotation, cell‒cell communication analysis, and functional annotation were used to investigate the impact of TAMs on the effectiveness of radiotherapy in NPC patients. Additionally, we employed RNA-seq data and clinical information from many patients, including 24 NPC samples from Zhujiang Hospital, 3,934 samples from patients with a history of radiotherapy, for further validation. We identified a hub TAM subpopulation, SPP1+ TAMs, that induced radioresistance in NPC based on single-cell RNA sequencing (scRNA-seq) data. We combined many public datasets to confirm their prognostic value in radiotherapy patients. We found that SPP1+ TAMs had stronger protumorigenic and immunosuppressive effects in the postradiotherapy recurrent tumor (PRRT) group. SPI1 transcription factor efficiently regulates the expression level of SPP1 in macrophages. In addition, we validated the underlying mechanisms by molecular docking, multiplex immunofluorescence and cellular experiments. In conclusions, by forming an immunosuppressive microenvironment and promoting tumor cell proliferation, SPP1+ TAMs leading to radioresistance and poor clinical prognosis in NPC patients. Targeting SPP1+ TAMs could provide new insights into improving the efficacy of radiotherapy in NPC patients.

## Background

Nasopharyngeal cancer (NPC) is a malignant tumor that arises in the nasopharynx and usually involves the posterior wall and the pharyngeal fossa. Owing to its concealed location, complex early symptoms, and vague characteristics, 70% of patients who are initially diagnosed with NPC are already in advanced stages [1]. Radiotherapy has significantly improved the treatment of NPC patients, and the five-year disease-specific survival (DSS) rate is 85% [2]. However, a subset of patients develop radioresistance after radiotherapy, resulting in poor clinical outcomes [3,4]. The increased DNA repair capacity of tumor cells is the major cause of radioresistances [5–7]. However, the precise details of the mechanism underlying radioresistance remain elusive.

The tumor microenvironment (TME) refers to the environment in which tumor cells originate and reside. It encompasses immune cells, stromal cells, tumor cells, and bioactive substances secreted by these cells [8]. Among the various immune cells found in tumor tissues, tumor-associated macrophages (TAMs) are particularly abundant [9]. TAMs can possess the ability to influence tumor proliferation, metastasis, and drug resistance phenotypes, rendering them a promising targets for antitumor therapy [9–11]. The function and phenotype of TAMs can influence the efficacy of radiotherapy. Research indicates that M1 TAMs enhance the effectiveness of radiotherapy, whereas M2 TAMs diminish it [12,13]. However, the M1 and M2 TAMs activation models do not fully represent the intricate in vivo environment [14,15]. Hence, further investigation is required to elucidate the specific role of different subsets of TAMs in conferring radioresistance in NPC.

This study aimed to comprehensively analyze the heterogeneity of TAMs and identify a TAM subpopulation, SPP1+ TAMs, that are critical for inducing resistance to radiotherapy in NPC based on scRNA-seq data. The prognostic significance of SPP1+ TAMs in radiotherapy patients has been confirmed in multiple public cohorts, and the potential mechanisms by which SPP1+ TAMs induce resistance to NPC radiotherapy have been elucidated by various techniques, including immunofluorescence, molecular docking, and cellular assays. This study provides novel insights into the development of personalized targeted therapy approaches for TAMs, as well as their combination with radiotherapy.

## Methods

This study included 15 NPC samples for single-cell RNA sequencing (scRNA-seq) and 24 NPC samples for RNA sequencing (RNA-seq). The patients were named the Zhujiang-NPC scRNA-seq and Zhujiang-NPC RNA-seq cohorts, and the two cohorts were completely independent, with no duplicate samples (sTable 1, 2). Patients were divided into a primary tumor (PT) group or a postradiotherapy recurrence tumor (PRRT) group. Tumor tissues in the PT group were derived from preradiotherapy, while tumor tissues in the PRRT group were derived from postradiotherapy recurrence. The inclusion and exclusion criteria are shown in appendix sTable 3.

Furthermore, clinical information and RNA-seq data of 3,934 patients with a history of radiotherapy, including pancancer (n=2,431), NPC (n=90), breast cancer (BRCA, n=1,137), prostate cancer (PRAD, n=248), head and neck cancer (HNSC, n=19), and locally advanced cervical cancer (LACC, n=9), were included in this study. Detailed information regarding all samples can be found in appendix sTable 4. The study protocol was approved by the Zhujiang Hospital academic committee (2019-KY-076-02). Informed consent was obtained before specimens were obtained from patients who underwent surgery at Zhujiang Hospital. Informed consent was waived for patients whose clinical information and RNA-seq data were obtained from publicly available datasets. This protocol has been registered at Chinese Clinical Trial Registry (identification number: ChiCTR2000028838).

### Single-cell suspension and RNA sample preparation

Detailed description of the process is provided in the supplement method.

### scRNA-seq and RNA-seq analysis

For the scRNA-seq data, fastq files and gene-barcode matrices were generated with 10X Cell Ranger software (version 7.0.1). The Seurat R package was used for quality control, analysis and visualization. To obtain valuable cells, several criteria were established: 1) < 200 genes/cell or > 8000 genes/cell and 2) > 10% of mitochondrial genes. After quality control, we obtained 120,579 cells. To remove batch effects, data integration was performed using canonical correlation analysis (CCA) and mutual nearest neighbor (MNN) algorithms. The top 2,000 variable features were identified for each sample, and integration was achieved using the “FindIntegrationAnchors” and “IntegrateData” functions. The FindClusters function was used to determine the number of cell clusters, and the resolution was set to 0.5. Cell-specific markers were used to annotate cell types. The FindMarkers function was used to identify differentially expressed genes (DEGs) between different subgroups, and the min.pct was set to 0.25. The screening criteria for DEGs were |logFC|>0.1 and P <0.05. The FindAllMarkers function was used to identify specific signature genes for different cell types. The top 50 DEGs for each cell type were considered cell-specific genes. The Startrac R package was used to calculate Ro/e values to assess cell tissue preferences. The CytoTRACE R package was used to assess cell differentiation potential, and the monocle3 R package was used to infer the developmental trajectory of cells. The AUCell R package was utilized for pathway quantification. The CellChat R package was used to determine cell‒cell communication patterns by inferring, analyzing and visualizing ligand‒receptor (LR) pairs between various cell types. Pyscenic software was used to construct a gene regulatory network analysis to identify gene regulatory networks based on the coexpression of gene regulators.

For the RNA-seq data, the raw sequencing reads were filtered and trimmed using Trim Galore with default parameters. Subsequently, they were mapped to the human genome GRCh38 using HISAT2. The calculation of gene expression levels was performed using FeatureCounts, and the counts were eventually converted to TPM values.

### Mechanism exploration

Gene set variation analysis (GSVA) and gene set enrichment analysis (GSEA) were used to determine the function of SPP1+ TAMs. The clusterProfiler R package was used for Gene Ontology (GO) and Kyoto Encyclopedia of Genes and Genomes (KEGG) analyses. For the scRNA-seq data, the AUCell R package was utilized for signature scoring. For bulk RNA-seq data, single-sample gene set enrichment analysis (ssGSEA) was utilized for signature scoring. The SPP1+ TAM level in a tumor estimate was estimated by the average expression level of CD68 and SPP1 [16]. The SPP1-CD44 LR pair in a tumor were used as the total gene expression values of SPP1and CD44 [17]. The related gene sets were obtained from MSigDb database. *P* < 0.05 was considered to indicate statistical significance. Detailed information on the signature gene sets used in this study is provided in appendix sTable 5.

### Immunofluorescence staining

After deparaffinizing the paraffin sections, performing antigen retrieval, and blocking with 3% H2O2 and 2% BSA, we sequentially applied different primary antibodies, including CD68, CD44 and SPP1, to the sections. The secondary antibody conjugated to horseradish peroxidase was then added dropwise and incubated for 50 minutes at room temperature. 4′,6-Diamidino-2-phenylindole dihydrochloride (DAPI) was used for nuclear staining of cells. Immunofluorescence microscopy was used to acquire and observe the images.

### Molecular docking

AlphaFold3 (https://golgi.sandbox.google.com/) was used to predict the structure of the new complex [18]. PDBePISA (https://www.ebi.ac.uk/msd-srv/prot_int/) was used to assess the binding energy and interfacial area of the complex [19].

### Experimental materials

Detailed description of the process is provided in the supplement method.

### Conditioned coculture and gene silencing

We inoculated THP-1 cells at a density of 1×105 cells/ml in a 6-well plate for 24 hours and then added 100 ng/ml PMA for 48 hours to treat the macrophage-like cells. Subsequently, we treated the cells with 1 mM tumor cell supernatant for 24 hours. To regulate our target gene, we used siRNA targeting SPI1 (SPI1-SI1 3’-5’: GCCCUAUGACACGGAUCUAUA SPI1-SI2 3’-5’: GAAGAAGCUCACCUACCAGUU)-transfected differentiated THP-1 cells, and all cells used for the experiments were free of mycoplasma or other prokaryotic contaminants.

### Real-time quantitative PCR (RT‒qPCR)

In this study, the primer sequences for SPI1 (Forward: GTGCCCTATGACACGGATCTA, Reverse: AGTCCCAGTAATGGTCGCTAT), SPP1 (Forward: GAAGTTTCGCAGACCTGACAT, Reverse: GTATGCACCATTCAACTCCTCG) and GAPDH (Forward: GGAGCGAGATCCCTCCAAAAT, Reverse: GGCTGTTGTCATACTTCTCATGG) were obtained from PrimerBank, and GAPDH was chosen as an internal reference to calculate the relative expression levels of SPI1 and SPP1 between different treatment groups via the 2-ΔΔCT method.

### Statistical analysis

All visualization and statistical analyses in this study were conducted using R software (version 4.1.2). The survminer R package was used to determine the optimal cutoff value. Kaplan‒Meier (KM) survival curves were used to present the differences in overall survival (OS), disease-free survival (DFS), progression-free survival (PFS), and DSS among different subgroups. A Cox regression model was utilized to calculate the hazard ratio (HR) and its corresponding 95% confidence interval (CI). For statistical tests of two groups, if the data had homogeneity in variance and passed the normality test, the parametric t test was used; otherwise, the nonparametric two-sample Wilcoxon rank test was used. Multivariate Cox regression analysis was conducted to identify independent prognostic factors. ROC curves were generated to determine the sensitivity and specificity of the selected features. All statistical analyses were two-tailed tests, and a significance level of *P* < 0.05 was adopted.

## Result

### Identifying the key role of TAMs in NPC radioresistance

To identify the predominant cell population contributing to radioresistance in NPC patients, 15 samples were collected for scRNA-seq; these included nine from the PRRT group and five from PT group. We also collected normal tissue from postradiotherapy recurrent adjacent tumor (PRRN). Thirteen major cell clusters were identified by representative marker genes (Figure 1A, B; Figure S1A, B), The proportion of TAMs was 6.67% (Figure 1C). To analyse differences in TME composition between the PRRN, PRRT and PT groups, the Ro/e values of various cell types were calculated (Figure S1C). The results showed a high level of infiltration of TAMs in the PRRT group. Differential expression analysis revealed that upregulated genes in the radioresistant group were mainly enriched in TAMs signature genes (Figure 1D-E). These results suggest that TAMs may be a crucial factor in radioresistance.

**Figure 1.**
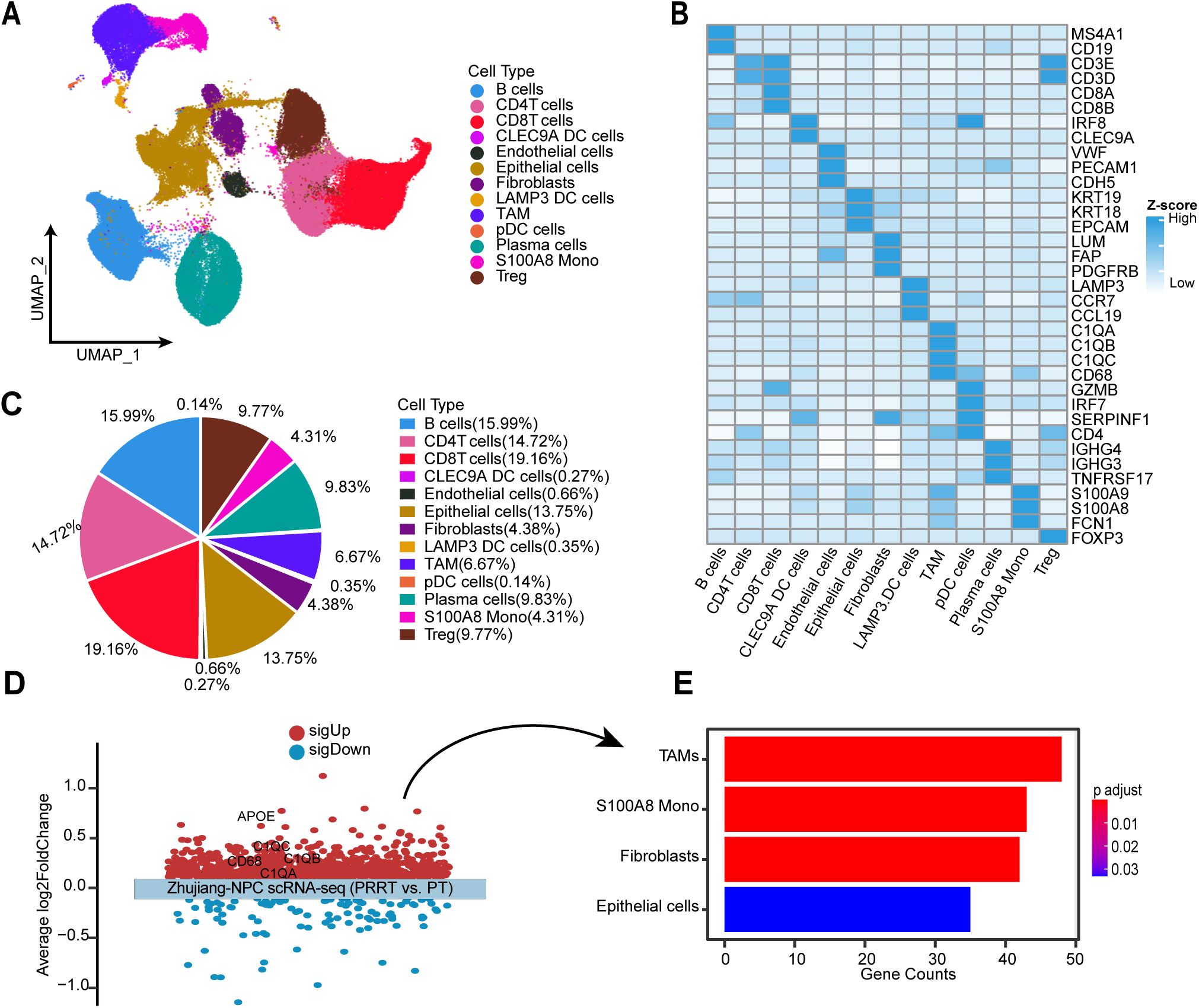
The landscape of the tumor microenvironment. (A) UMAP projection of major cell types coloured by cell type. (B) Heatmap showing representative markers of major cell types. (C) Pie plot showing the proportions of major cell types. (D) Scatter plot of differential expression analysis between the PRRT and PT groups. (E) Bar plot showing enrichment analysis of upregulated genes in the PRRT group. PRRT, post-RT recurrent tumor; PT, primary tumor; TAMs, tumor-associated macrophages.

### Identification and characterization of TAM diversity

To further characterize TAMs, three subclusters (named TAM-1, TAM-2 and TAM-3) were identified and visualized using UMAP plot (Figure 2A, Figure S2A). ROGUE analysis revealed good homogeneity of all three TAM subpopulations across samples (Figure 2D). The TAM-1 cluster was significantly enriched in the PRRT group, and TAM-3 cluster was significantly enriched in the PT group (Figure 2C). The proportion of TAMs-1 was 34.4% (Figure 2B). The heatmap revealed the top 10 marker genes for each subcluster, with APOE and SPP1 being highly expressed in the TAM-1 cluster, CXCL10 and IL1B in the TAM-2 cluster, and S100A8 and S100A9 in the TAM-3 cluster (Figure 2E, sTable 6). To explore the function of each subcluster, enrichment analysis was performed using their signature genes. The results indicated that TAM-1 cluster was enriched in lipid metabolism, leukocyte migration, immune system, and complement activation pathways. In contrast, TAM-2 cluster was enriched in leukocyte chemotaxis and migration-related pathways, and TAM-3 cluster was enriched in phagocytosis, bacterial defence, immune system, and complement activation pathways (Figure S2B-D). These results indicate that TAMs exhibit high heterogeneity.

**Figure 2.**
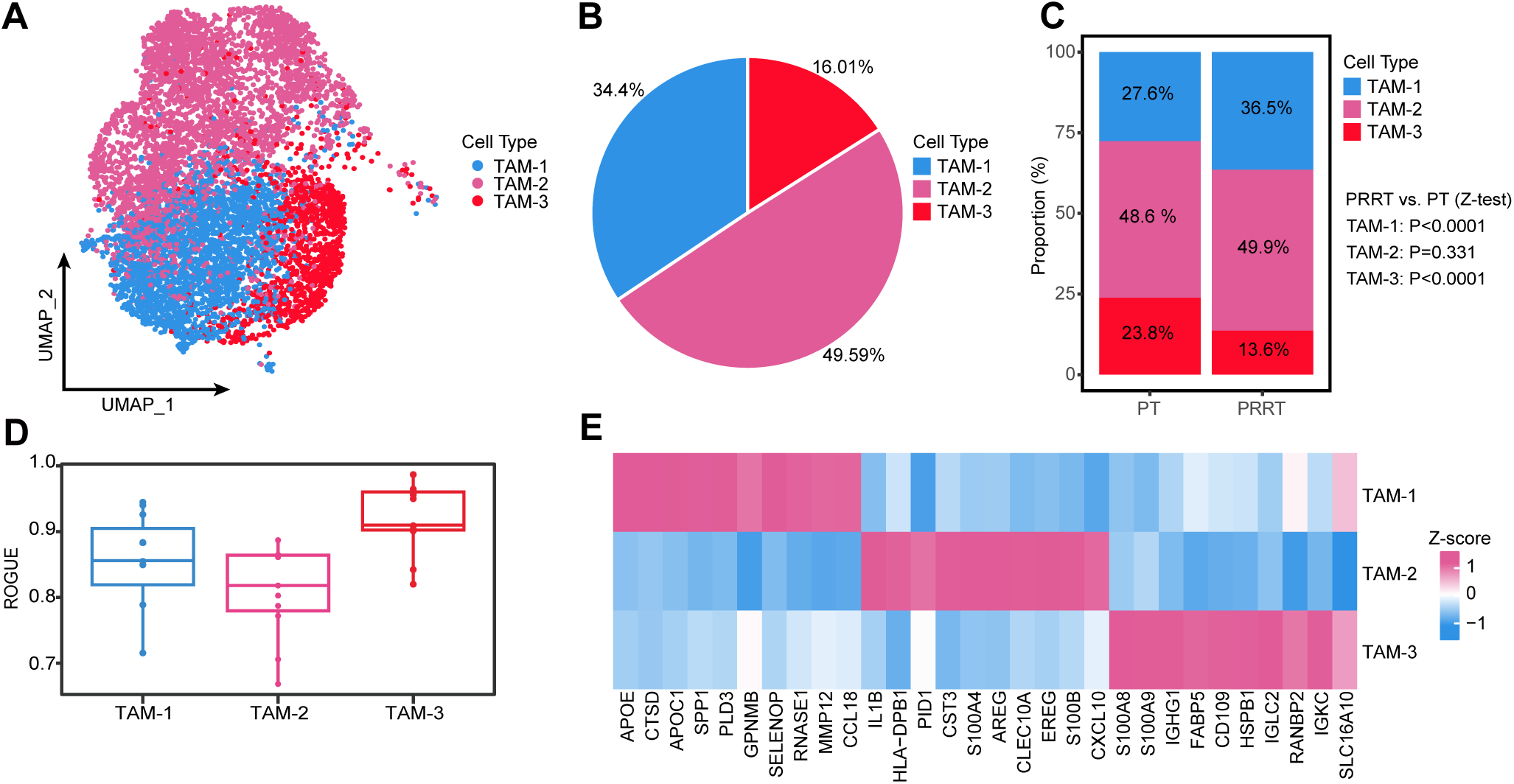
The heterogeneity of TAMs. (A) UMAP projection of TAMs coloured by subcluster. (B) Pie plot showing the proportions of TAM subclusters. (C) Bar plot showing the proportions of TAM subclusters in the PRRT and PT groups. (D) Evaluation of the purity of TAM subclusters. (E) Heatmap showing the top 10 specific signature genes of various TAM subclusters. PRRT, postradiotherapy recurrent tumor; PT, primary tumor; TAMs, tumor-associated macrophages.

### Identification the clinical value of SPP1+ TAMs

To identify the key TAM subclusters in radioresistance, the Ro/e values were calculated for each TAM subcluster. The heatmap also showed that the TAM-1 cluster was enriched in the PRRT group (Figure 3A). The dot plot shows that SPP1 was highly expressed specifically in the TAM-1 cluster (Figure 3B). In the Zhujiang-NPC cohort, there was grater infiltration of SPP1+ TAMs in the PRRT group than in the PT group and the number of SPP1+ TAMs was effective at predicting the sensitivity of patients to radiotherapy (Figure 3C, D; Figure S3). Immunofluorescence also revealed significant enrichment of SPP1+ TAMs in the PRRT group. KM survival analysis revealed a significant association between increased infiltration of SPP1+ TAMs and unfavourable OS, PFS, DFS, and DSS in radiotherapy patients (Figure 4A; Figure S4A-C). The three independent datasets also showed the same trend (Figure 4B-D).

**Figure 3.**
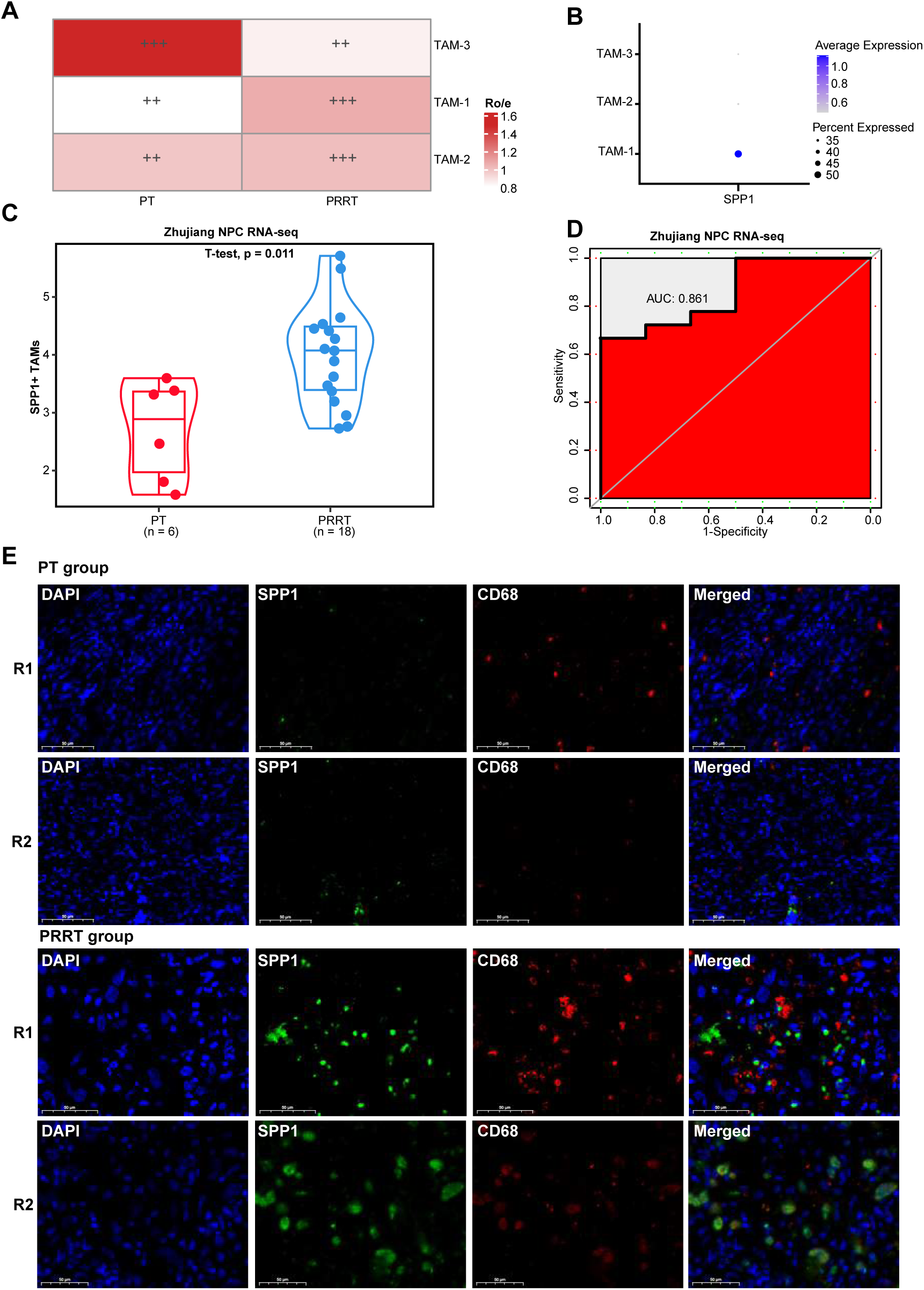
The clinical value of SPP1+ TAMs. (A) Tissue preference of each major cell type estimated by the Ro/e value. +++, Ro/e > 1; ++, 0.8 < Ro/e ≤ 1; +, 0.2 ≤ Ro/e ≤ 0.8; +/−, 0 < Ro/e < 0.2; −, Ro/e = 0. (B) Dot plot showing SPP1 expression in TAMs subclusters. (C) Box plot showing the infiltration of SPP1+ TAMs in the PRRT and PT groups. (D) ROC curves showing the predictive value of the number of SPP1+ TAMs for patients receiving radiotherapy. (E) Immunofluorescence staining of CD68 and SPP1 in NPC samples from the Zhujiang-NPC RNA-seq cohort showing the infiltration of SPP1+ TAMs in the PRRT and PT groups. PRRT, postradiotherapy recurrent tumor; PT, primary tumor; TAMs, tumor-associated macrophages.

**Figure 4.**
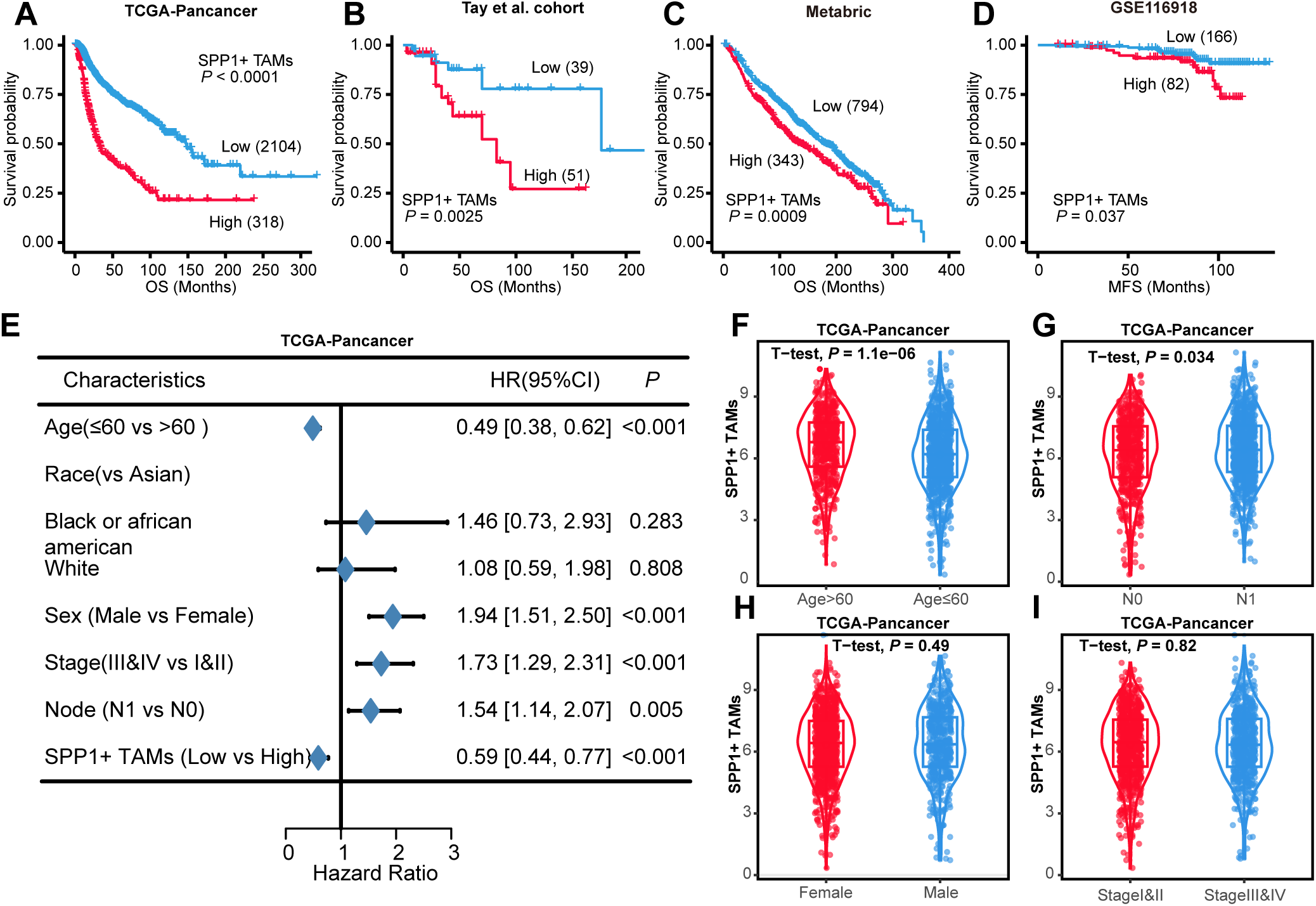
Association of SPP1+ TAMs with clinical features. (A-D) Survival curves demonstrating survival differences between the high– and low-infiltration SPP1+ TAM groups. (E) Forest plot demonstrating multivariate analysis of SPP1+ TAMs and clinicopathological features. (F-G) Box plot revealing the relationship between the infiltration level of SPP1+ TAMs and age, stage, sex, and lymph node status. TAMs, tumor-associated macrophages.

To validate whether the number of SPP1+ TAMs is an independent prognostic factor for patients receiving radiotherapy, multivariate analysis was conducted. We identified certain clinical parameters as independent prognostic factors, including age, sex, stage and lymph node status (Figure 4E). The infiltration of SPP1+ TAMs was significantly correlated with age and lymph node status but not with sex or stage (Figure 4F-I). In addition, the KM survival curves showed that patients with a low infiltration of SPP1+ TAMs had a better OS than those with a high infiltration of SPP1+ TAMs in the same clinical subgroup (Figure S5A-C).

### Identification and characterization of cell fate transitions and functional profiles of SPP1+ TAMs

To investigate the cell fate transition of SPP1+ TAMs, Monocle3 was used to infer TAM trajectories. CytoTRACE and Monocle3 analyses indicated that TAM-1 originated from TAM-2, which had the highest CytoTRACE score and lowest pseudotime value, and developed a termediated state of TAM-3 (Figure 5A-D). SPP1 expression increased along the trajectory of TAM transition (Figure 5E, F). Energy metabolism pathway analysis revealed that glycolysis, hypoxia, glutamine metabolism and lipid metabolism pathway activities were significantly greater in SPP1+ TAMs than in SPP1-TAMs (Figure 5G-J). The GSEA results indicated a significant enrichment of signature genes of hypoxia, angiogenesis, and NFKB pathways in SPP1+ TAMs (Figure 5K). Consistent results were obtained from four independent datasets (Figure S6C). Moreover, the expression of SPP1 was greater in M2 TAMs, and the M2 signature score was greater in SPP1+ TAMs (Figure S1 A, B). These results imply that SPP1+ TAMs might constitute an M2 TAM subset associated with proangiogenic and immunosuppressive functions.

**Figure 5.**
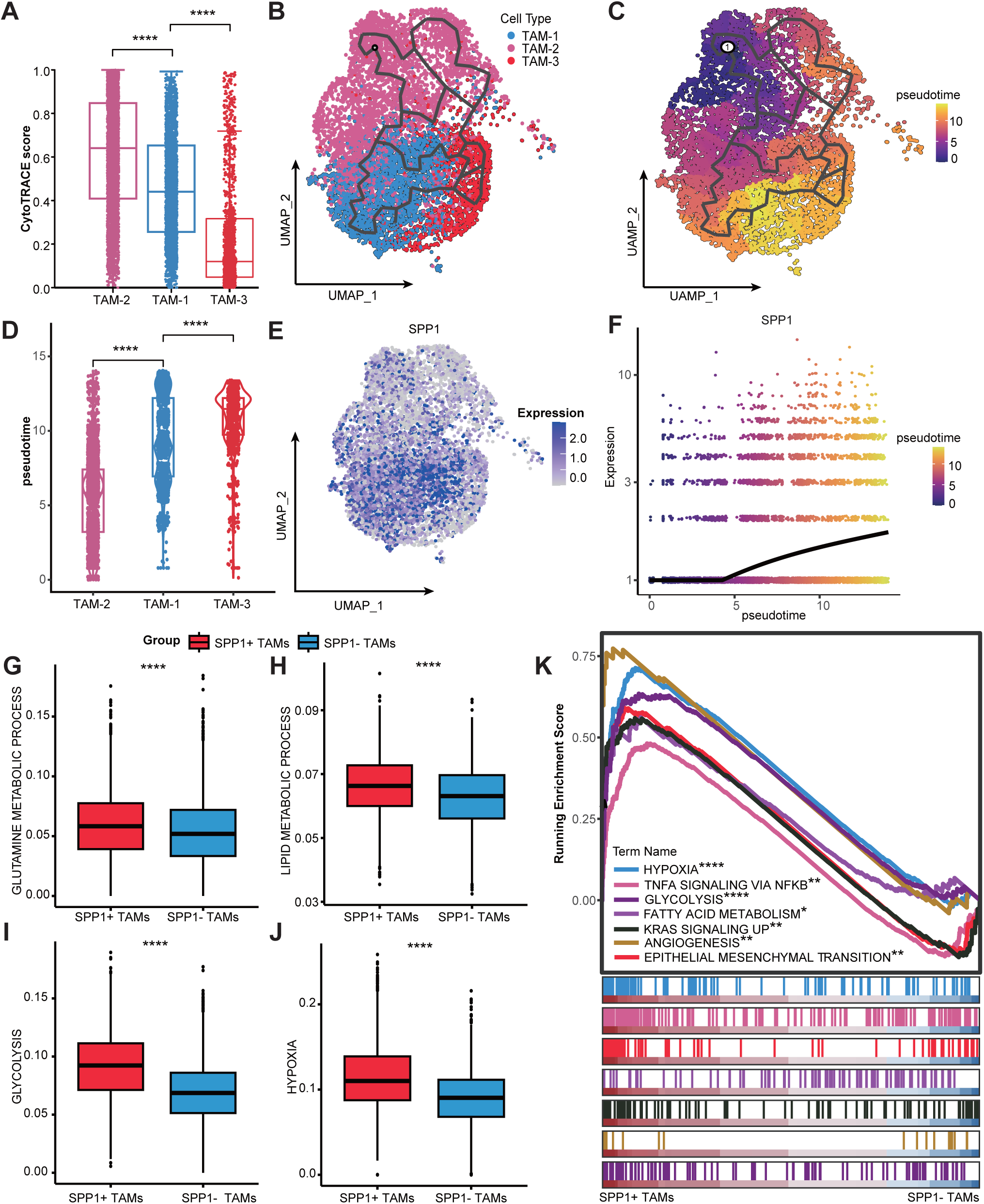
The functional profiles and differentiation status of SPP1+ TAMs. (A) Boxplots demonstrating the difference in CytoTRACE score among TAM-1, TAM-2 and TAM-3 clusters. Higher CytoTRACE score means higher differentiation potential. (B-C) UMAP plot showing the pseudotime trajectories of TAMs. (D) Boxplots demonstrating the differences in pseudotime values among TAM-1, TAM-2 and TAM-3 clusters. (E) UMAP plot exhibiting the distribution of SPP1 in TAMs. (F) Relationship between SPP1 expression and pseudotime values. (G-J) Boxplot demonstrating differences in glycolysis, hypoxia, lipid metabolism, and glutamine metabolism pathway activity scores between SPP1+ TAMs and SPP1-TAMs. (K) GSEA between the SPP1+ TAMs and SPP1-TAMs evaluated in the context of hallmark gene sets. TAMs, tumor-associated macrophages. *P < 0.05; **P < 0.01; ***P < 0.001; ****P < 0.0001.

### Identification of the interaction network of SPP1+ TAMs

To decode cell‒cell communication in the NPC, CellChat was used to explore potential LR pairs between SPP1+ TAMs, epithelial cells, endothelial cells and T cells. The intensity and number of cell‒cell communications were greater in the PT group than in the PRRT group (Figure S7A, B). Enrichment analysis revealed that SPP1+ TAMs communicated most frequently through LR pairs in the VEGF, CD86 and CD80 pathways for cell ‒ cell communication (Figure 6A, B; Figure S8A). A comparison of overall communication probabilities revealed that thirteen pathways, including the VEGF and complement pathways, were highly active in the PRRT group, and ten pathways, including the MHC-I and CXCL pathways, were highly active in the PT group (Figure 6C, Figure S8B-E). The interaction between SPP1+ TAMs and T cells (CD4, CD8 T cells and Treg cells) were mainly related to immunosuppression-related ligands and receptors, such as TNF:TNFRSF1B, NECTIN2:TIGIT, and LGALS9:HAVCR2 (Figure 6C, D). SPP1+ TAMs can also inhibit T-cell function by interacting with regulatory T cells (Tregs) through the CD80/CD86-CTLA4 axis (Figure 6E; Figure S8F). SPP1+ TAMs interacted with epithelial and endothelial cells via LR pairs on the EGFR and VEGF pathways, promoting tumor cell proliferation (Figure 6 H, I). In addition, we observed that the SPP1-CD44 LR pair mediated cell‒cell communication between SPP1+ TAMs and various cell types (Figure 6C-G). Multiple immunofluorescence assays consistently demonstrated the proximity of SPP1+ TAMs to CD44+ cells (Figure 6H). KM survival curves showed that radiotherapy patients with high expression of the SPP1-CD44 LR pair had a poor prognosis (Figure 6I; Figure S7C). Interestingly, the pro-angiogenic and immunosuppressive functions of SPP1+ TAMs were stronger in the PRRT group than in the PT group (Figure S8G, H). These findings emphasized that SPP1+ TAMs promote TME remodelling and immunosuppression in NPC.

**Figure 6.**
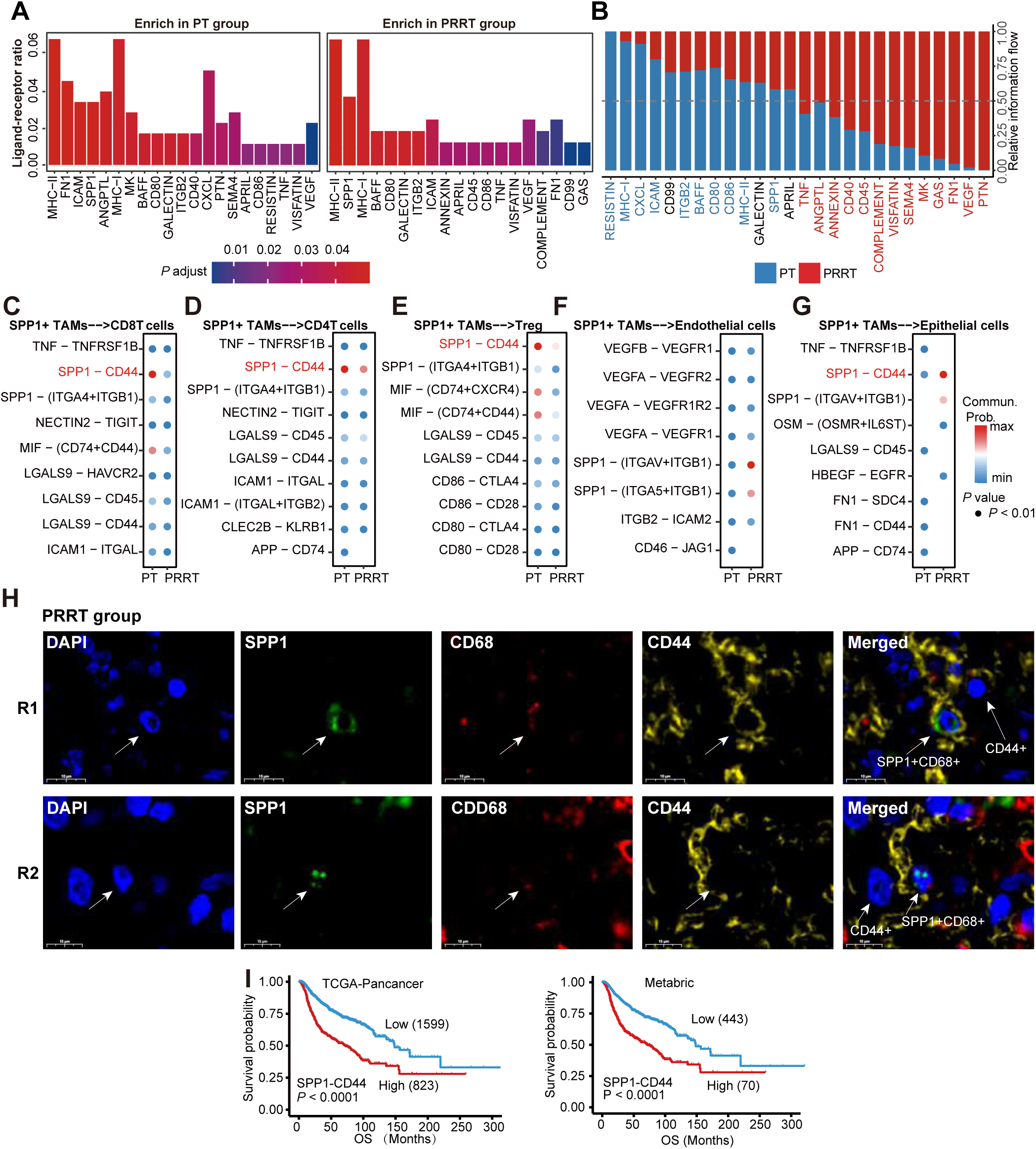
The cell‒cell communication of SPP1+ TAMs. (A) Bar plot showing the enrichment analysis of SPP1+ TAMs LR pairs in the PRRT and PT groups. (B) Bar plot showing the selected pathways ranked based on their differences in overall information flow within the inferred networks between the PRRT and PT groups. The red pathways are significantly enriched in the PRRT group. The black pathways indicates equal enrichment in the PRRT and PT groups. The blue pathways significantly enriched in the PT group are shown. (C-G) Dot plot demonstrating LR pairs of SPP1+ TAMs interacting with CD8 T cells, CD4 T cells, Tregs, endothelial cells and epithelial cells. (H) Immunofluorescence staining of CD44 and SPP1 in NPC samples from the Zhujiang-NPC RNA-seq cohort. (I) The prognostic value of SPP1–CD44 ligand–receptor pairs. NPC, nasopharyngeal cancer; PRRT, postradiotherapy recurrent tumor; PT, primary tumor; TAMs, tumor-associated macrophages.

### Identification of specific transcription factors (TFs) of SPP1+ TAMs

To identify TFs specific to SPP1+TAMs, a pyscenic software was conducted. We identified six major modules in myeloid cells, with module 5 exhibiting the highest activity in SPP1+ TAMs (Figure 7A, B; Figure S9A). Eight transcription factors are risk factors for three or four clinical indicators in radiotherapy patients, seven of which were significantly upregulated in SPP1+ TAMs (Figure 7C, D). To further investigate the TFs in SPP1+ TAMs, their specificity score was assessed using Jensen–Shannon divergence [20]. SPI1 had the highest specificity score (Figure 7E). In addition, alphafold3 and PDBePIA analyses showed that the SPI1 transcription factor and the promoter of SPP1 can form a stable complex. (Figure 7F). Supernatants from tumor cells were able to cause significant upregulation of SPP1 in human macrophages (Figure 7G). In contrast, silencing SPI1 with siRNA significantly reduced SPP1 expression. Functional analysis revealed that the target genes of SPI1 were enriched in the DNA repair, oxidative phosphorylation and MYC target V1 pathways (Figure S9B). These results emphasize the important role of SPI1 in the development of SPP1+ TAMs.

**Figure 7.**
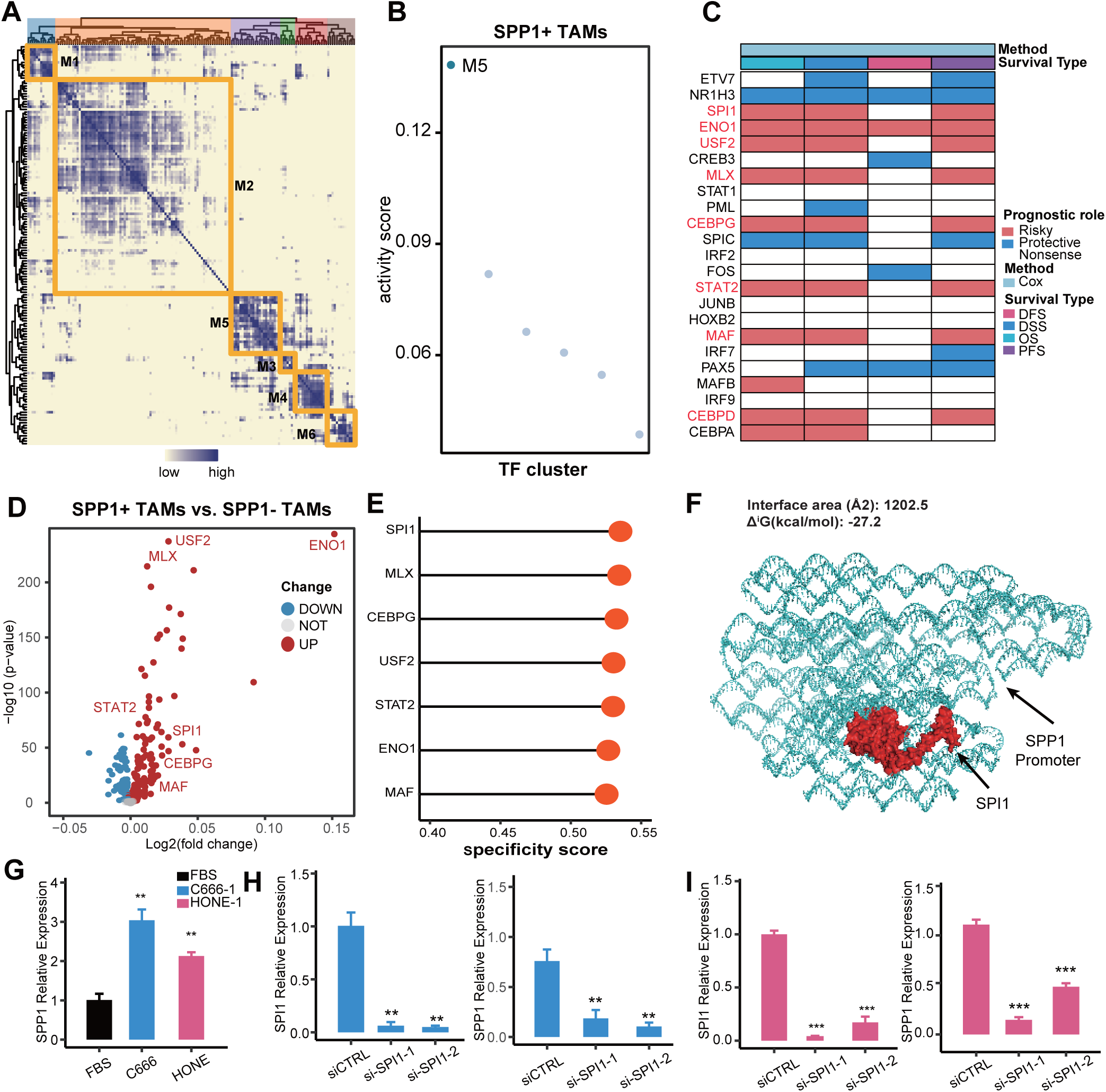
Specific transcription factors (TFs) of SPP1+ TAMs. (A) Heatmap showing TFs clustering. (B) Scatter plot demonstrating the activity scores of TF clusters. (C) Heatmap demonstrating the predictive value of the M5 cluster of TFs for clinical outcomes in the TCGA-Pancancer cohort. (D) Volcano plot of TFs between SPP1+ TAMs and SPP1-TAMs. (E) Specificity scores of the selected TFs for SPP1+ TAMs. (F) The new complex between the SPI1 protein and the SPP1 promoter. (G) PMA-induced THP-1 cells were treated with supernatants from C666 and HONE cells. (H, I) PMA-induced THP-1 cells were treated with siRNA and supernatants from C666 (H) or HONE (I) cells. TAMs, tumor-associated macrophages; TFs, transcription factors. *P < 0.05; **P < 0.01; ***P < 0.001; ****P < 0.0001.

## Discussion

Radiotherapy is the primary treatment modality for patients with NPC [21, 22]. However, the development of radioresistance significantly limits the effectiveness of radiotherapy. Numerous studies have demonstrated a close association between the development of radioresistance and TAMs [19–21]. The contribution of TAM heterogeneity to radioresistance in NPC has not been thoroughly investigated. This study involved the collection and analysis of scRNA-Seq data from 15 patients with NPC at Zhujiang Hospital and identified a distinct subset of M2 TAMs characterized by high SPP1 expression and a mid-to-late phenotype. Functional analysis indicated that SPP1+ TAMs exhibited a proangiogenic and immunosuppressive phenotype and were significantly enriched in the PRRT group. The infiltration level of SPP1+ TAMs could serve as an effective predictor of radiotherapy sensitivity and clinical outcomes in patients with NPC.

TAMs are among the most abundant immune cells present in tumor tissues, and the number of TAMs can serve as a marker for predicting the efficacy of radiotherapy across various types of tumors [26]. The heterogeneity of TME is closely associated with the effectiveness of clinical treatment [27–29]. CD163+ TAMs are significantly associated with poor clinical outcomes in radiotherapy patients with head and neck cancer or chondroblastoma [30–31]. In the present study, SPP1+ TAMs mediated NPC radioresistance through their protumorigenic and immunosuppressive functions, which are associated with poor prognosis in radiotherapy patients.

The function and phenotype of TAMs are intricately linked to the effectiveness of radiotherapy [12,13]. Research has demonstrated that M1 TAMs can bolster the effectiveness of radiotherapy through the inhibition of proangiogenic cytokines, including IL-12 and TNFα, as well as by triggering T-cell-mediated antitumor immune responses [32]. Conversely, M2 TAMs can decrease the effectiveness of radiotherapy by upregulating VEGF via the TNFα/TNFR signalling pathway [33]. In this study, SPP1 expression was increased along the trajectory of TAM transition, which can promote macrophage polarization toward the M2 phenotype [34]. Moreover, functional analysis demonstrated that SPP1+ TAMs exhibited significant enrichment in hypoxia, NFKB, and angiogenesis signalling pathways compared to SPP1-TAMs. The NFKB pathway diminishes the effects of radiotherapy by attenuating the DNA damage caused by irradiation [31]. Hypoxia not only promotes tumor cell proliferation but also suppresses immune function by creating an acidic environment [36, 37]. SPI1 is recognized as a key TF in macrophage development [38]. Our findings further suggest that SPI1 is specifically expressed in SPP1+ TAMs and may promote the formation of protumorigenic and immunosuppressive functional phenotypes in SPP1+ TAMs. These findings suggest that SPP1+ TAMs exhibit an M2 phenotype, consistent with previous studies reporting their protumorigenic and immunosuppressive functions [34]. Interestingly, the protumorigenic and immunosuppressive functions of SPP1+ TAMs were stronger in the PRRT group, which may be related to the specific TME of patients in the PRRT group.

In the TME, intricate cell‒cell interactions occur among diverse cell types. These interactions not only facilitate tumor cell growth but also play a vital role in promoting tumor cell adaptability and growth despite treatment [39, 40]. M2 TAMs promote tumor progression by promoting the formation of an immunosuppressive microenvironment and angiogenesis [41, 42]. In the present study, SPP1+ TAMs can inhibit T-cell function, which have been shown to be a risk factor in multiple tumors [38–41]. Specifically, SPP1+ TAMs can inhibited T-cell-mediated antitumor immunity by utilizing multiple immunosuppressive LR pairs and interacting with Treg cells through the CD80/CD86-CTLA4 axis. Moreover, SPP1+ TAMs can stimulate tumor vascular proliferation and tumor cell growth by interacting with endothelial and epithelial cells via LR pairs on the VEGF and EGFR pathways. In addition, previous studies have shown that the SPP1-CD44 interaction promotes tumor progression [47]. Our findings suggest that this interaction is involved in the development of radioresistance in NPC. These findings highlight the significant contribution of SPP1+ TAMs to the development of radioresistance in NPC.

This study has several limitations that should be addressed. First, additional in vivo and in vitro functional validation of SPP1+ TAMs is needed. Second, the sample size of local NPC patients included in this study was only 24 patients. To supplement the validation analysis, pancancer patients with a history of radiotherapy were also included. However, this may introduce heterogeneity among different cancer types. Nevertheless, these findings suggest the potential role of SPP1+ TAMs in inducing radioresistance across various cancers, which warrants further investigation.

## Conclusion

In conclusion, SPP1+ TAMs are a subset of TAMs with pro-angiogenesis and immunosuppressive phenotypes. These TAMs can reshape the immune microenvironment and form an immunosuppressive microenvironment, leading to radioresistance and poor clinical prognosis in NPC patients.Targeting SPP1+ TAMs could provide new insights into improving the efficacy of radiotherapy in NPC patients.

## Abbreviations

CI: confidence interval
DSS: disease-specific survival
DEGs: differentially expressed genes
DFS: disease-free survival
GSVA: Gene set variation analysis
GSEA: gene set enrichment analysis
HR: hazard ratio
KM: Kaplan‒Meier
NPC: nasopharyngeal carcinoma
OS: overall survival
PFS: progression-free survival
PRRT: post-radiotherapy recurrent tumor
PT: primary tumor
RNA-seq: RNA sequencing
scRNA-seq: single-cell RNA sequencing
TME: tumor microenvironment
TAMs: tumor-associated macrophages.

## Acknowledgements

We thank the patients, their families and those who participated in the study. In addition, We thank Dr.Jianming Zeng(University of Macau), and all the members of his bioinformatics team, biotrainee, for generously sharing their experience and codes; as well as all members who build and maintain TCGA and GEO databases

## Authors’ contributions

Conceptualization, J.Z., P.L., and H.M.; Formal analysis, W.D.Z., L.L.L., H.X.Y., X.Y.Z., Q.Y., N.L.; Software, W.D.Z., L.L.L.; Supervision, J.Z., P.L., and H.M.; Visualization, W.D.Z., N.L., L.L.L., H.X.Y., J.Y.C., S.L.H., Q.C., Z.Q.L.; Writing–original draft, W.D.Z., L.L.L., X.Y.Z., A.Q.L; Writing–review & editing, W.D.Z., L.L.L., X.H.Y., X.Y.Z., S.L.H., J.Z., H.M., and P.L. All authors read and approved the final manuscript.

## Availability of data and materials

All data of this study are available within the article and the supplemental information and from the corresponding author upon reasonable request.

## Declarations

### Ethics approval and consent to participate

The study’s protocol was approved by the Zhujiang Hospital academic committee (2019-KY-076-02). Informed consent was obtained before obtaining specimens from patients undergoing surgery at Zhujiang Hospital. Informed consent was waived for patients whose clinical information and RNA-seq data were obtained from publicly available datasets. This protocol has been recistered at Chinese Clinical Trial Registry (identification number: ChiCTR2000028838).

### Consent for publication

All authors consent this manuscript to be published.

